# Enhanced neutralization escape to therapeutic monoclonal antibodies by SARS-CoV-2 Omicron sub-lineages

**DOI:** 10.1101/2022.12.22.521201

**Authors:** Franck Touret, Emilie Giraud, Jérôme Bourret, Flora Donati, Jaouen Tran-Rajau, Jeanne Chiaravalli, Frédéric Lemoine, Fabrice Agou, Etienne Simon-Lorière, Sylvie van der Werf, Xavier de Lamballerie

## Abstract

The landscape of SARS-CoV-2 variants dramatically diversified with the simultaneous appearance of multiple sub-variants originating from BA.2, BA.4 and BA.5 Omicron sub-lineages. They harbor a specific set of mutations in the spike that can make them more evasive to therapeutic monoclonal antibodies. In this study, we compared the neutralizing potential of monoclonal antibodies against the Omicron BA.2.75.2, BQ.1, BQ.1.1 and XBB variants, with a pre-Omicron Delta variant as a reference. Sotrovimab retains some activity against BA.2.75.2, BQ.1 and XBB as it did against BA.2/BA.5, but is less active against BQ.1.1. Within the Evusheld/AZD7442 cocktail, Cilgavimab lost all activity against all subvariants studied, resulting in loss of Evusheld activity. Finally, Bebtelovimab, while still active against BA.2.75, also lost all neutralizing activity against BQ.1, BQ.1.1 and XBB variants.

## Main

Since the emergence of the Severe Acute Respiratory Syndrome Coronavirus 2 (SARS-CoV-2) in China in late 2019, vaccines have been the most effective and widely used therapy. However, a fraction of the population does not respond to immunization (*i*.*e*, immuno-compromised). There monoclonal antibodies (mAbs) have proven a great resource, both for prevention and treatment of infection^1^. Most of such mAbs have been developed during the early stages of the outbreak and target the original SARS-Cov-2 spike^1,2^. One of them was developed from a SRAS survivor and is a broadly neutralizing antibody^3^. Unfortunately, the most recent circulation of SARS-CoV-2 has been associated with the spread of multiple sublineages (*i*.*e*., Omicron BA.1, BA.2, BA.4, BA.5 and more recently BA.2.75.2, BQ.1, BQ.1.1 and XBB variants) that combine increased transmissibility and immune escape^4-9^. They harbor different mutations in the spike that can make them more evasive to vaccination and infection induced antibodies as well as therapeutic monoclonal antibodies^10^.

Specifically, BA.2.75.2 is derived from BA.2 and contains, among others, three major additional mutations in the Receptor-Binding Domain (RBD): R346T, N460K and F486S (Fig 1, Supp fig 1,3) among which N460K and F486S are located in the ACE-2 Receptor Binding Motif (RBM). BQ.1 and BQ.1.1 are direct descendants from BA.5 and therefore contain the F486V mutation^8^. BQ.1 has gained K444T and N460K while BQ.1.1 has in addition the R346T mutation (Fig 1, Supp fig 1,3). Finally, the XBB variant is the result of a single breakpoint recombination in the RBD between two BA.2 sub-variants: BJ.1 which contains the R346T mutation and BM.1.1.1 derived from BA.2.75 with the addition of the F486S and F490S mutations (Fig 1, Supp fig 1,3).

**Figure 1:**
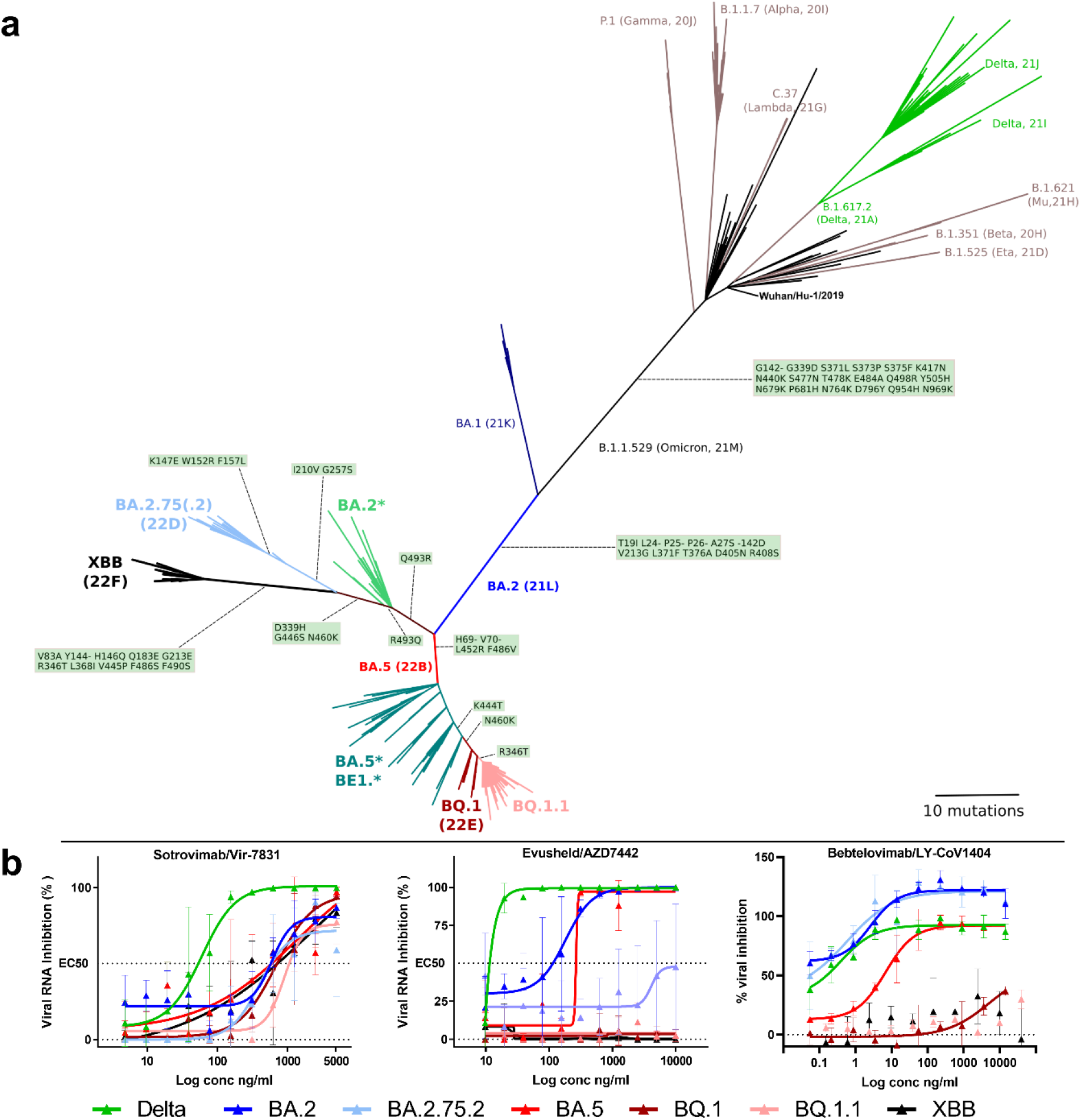
Omicron subvariants phylogenetic analysis and susceptibility to therapeutic monoclonal antibodies. a) Unrooted phylogenetic tree displaying BQ.1.1 and XBB lineages in the context of SARS-CoV-2 main lineages; amino-acid mutations in the Spike are displayed on branches of the tree for lineages of interest. The complete set of amino-acid mutations is depicted in Supp. fig 3. b) dose response curves reporting the susceptibility of the SARS-CoV-2 Delta pre-omicron variant and Omicron subvariants to a panel of therapeutic monoclonal antibodies. Antibodies tested: Sotrovimab/Vir-7831, Evusheld/AZD7742 cocktail and Bebtelovimab/Ly-CoV1404. Data presented are from three technical replicates in VeroE6-TMPRSS2 cells, and error bars show mean±s.d.

The appearance of recurrent mutations or mutations at recurrent positions in different sublineages suggests a convergent evolution of the Omicron RBD as a result of humoral immunity to SARS-CoV-2 in the population^10,11^.

After three successive waves of Omicron (BA.1, BA.2, and BA.4/BA.5) only few therapeutic monoclonal antibodies neutralizing these variants remained active^5,12,13^. The convergent RBD mutations observed in the multiple Omicron sub-variants have the potential to further alter the activity of available therapeutic monoclonal antibodies.

In this study, we tested the neutralizing activity of therapeutic antibodies against clinical isolates of the BA.2.75.2, XBB, BQ.1 and BQ.1.1 sub lineages. We used different sets of clinical isolates as control; for BA.2.75.2 and XBB we used their first progenitor BA.2 and similarly we used BA.5 for BQ.1 and BQ.1.1. The Delta pre-Omicron variant (lineage B.1.617.2) was used as reference for antibody neutralizing activity^14^.

We tested therapeutic antibodies currently in use that have been shown to retain neutralizing activity against previous Omicron sub-variants^13^ along with the Roche Regeneron antibodies Casirivimab (REGN10933) and Imdevimab (REGN10987), which regained activity against BA.2^15^.

All these monoclonal antibodies target the spike Receptor Binding Domain (RBD)^2,16^. However, based on analysis of their structure in complex with the RBD showing that they exhibit different binding modes, they were classified into two distinct anti-RBD antibody classes^17^. Sotrovimab/Vir-7831, which is derived from parental antibody S309, and belongs to class 3 neutralizing antibodies, has been isolated and developed from a SARS-CoV survivor and targets the RBD core region, outside the RBM^3^. Like Sotrovimab, Cilgavimab/AZD1061, Imdevimab (REGN10987) and Bebtelovimab (LY-Cov1404) belong to the same structural class and bind outside the RBM^2,16,18,19^. Finally Tixagevimab/AZD8895 and Casirivimab/REGN10933 are targeting the RBM^2,16,18^, and belong to the class 1 Nabs.

We applied a standardized protocol for the evaluation of antiviral compounds based on the reduction of RNA yield^20,21^, which has been applied previously to SARS-CoV-2 antivirals and therapeutic antibodies evaluation^13,22,23^. This assay, based on authentic and replicating viruses, was performed in VeroE6 TMPRSS2 cells; the viral RNA in the supernatant medium was quantified by qRT-PCR at 48h post-infection to determine the 50% effective concentration (EC_50_).

We first observed a complete loss of detectable neutralizing activity for the four sub-variants with Imdevimab (REGN10987) (Table 1, Supp Fig.2), and still no activity with Casirivimab which made it impossible to calculate the EC50 (Table 1, Supp Fig.2). This result is in line with previous reports using a pseudo-virus assay^10,24^, live virus^25^ and with a study using a fusogenicity reporter assay^26^.

**Table 1:** Activity of therapeutic antibodies against Delta and Omicron BA.2, BA.5, BA.2.75.2, BQ.1 and BQ.1.1 variants. Interpolated EC_50_ values are expressed in ng/mL. Sotrovimab EC_50_ values for Delta, BA.2.75.2, BQ.1, BQ1.1 and XBB are the mean of two independent experiments. For the control strain Delta, EC_50_ is the mean of two independent experiments (n.n: non-neutralizing). Fold change reductions were calculated in comparison with the pre-Omicron Delta strain.

|  |  |  | Strain |  |  |  |  |  |  |
| --- | --- | --- | --- | --- | --- | --- | --- | --- | --- |
| Antibody |  |  | Delta | BA.2 | BA.2.75.2 | BA.5 | BQ.1 | BQ.1.1 | XBB |
| Regeneron | Casirivimab<br>REGN10933 | EC<sub>50</sub><br>fold-change | 14.7<br>- | n.n<br>- | n.n<br>- | n.n<br>- | n.n<br>- | n.n<br>- | n.n<br>- |
|  | Imdevimab<br>REGN10987 | EC<sub>50</sub><br>fold-change | 20.1<br>- | 345.9<br>17.2 | n.n<br>- | 254.4<br>12.7 | n.n<br>- | n.n<br>- | n.n<br>- |
| GSK/ Vir | Sotrovimab<br>(vir-7831) | EC<sub>50</sub><br>fold-change | 98.6<br>- | 578.9<br>5.9 | 422.3<br>4.3 | 596.5<br>6.0 | 688.2<br>7.0 | 1238.0<br>12.6 | 806.6<br>8.2 |
| AstraZeneca | Cilgavimab<br>(AZD1061) | EC<sub>50</sub><br>fold-change | 21.7<br>- | 184.0<br>8.5 | n.n<br>- | 147.4<br>6.8 | n.n<br>- | n.n<br>- | n.n<br>- |
|  | Tixagevimab<br>(AZD8895) | EC<sub>50</sub><br>fold-change | 12.4<br>- | n.n<br>- | n.n<br>- | n.n<br>- | n.n<br>- | n.n<br>- | n.n<br>- |
|  | Evusheld<br>(AZD7442) | EC<sub>50</sub><br>fold-change | 12.5<br>- | 114.9<br>9.2 | n.n<br>- | 265.4<br>21.2 | n.n<br>- | n.n<br>- | n.n<br>- |
| Lilly | Bebtelovimab<br>(LY-Cov1404) | EC<sub>50</sub><br>fold-change | 0.4<br>- | 2.6<br>6.7 | 0.6<br>1.5 | 6.8<br>17.4 | n.n<br>- | n.n<br>- | n.n<br>- |

n.n, non-neutralizing at the highest concentration tested

Our results show that Sotrovimab retains some neutralizing activity against BA.2.75.2, XBB, BQ.1 an BQ.1.1 *in vitro*^10,24,26^. Regarding BA.2.75.2, Sotrovimab activity seems unchanged compared to its progenitor BA.2 and a modest decrease in neutralization is observed with the XBB variant (estimated EC_50_ of 806.6ng/mL). In the case of the BQ.1 variant (Table 1, Fig.1), the EC_50_ increases modestly from 596.5 (BA.5) to 688.2 (BQ.1) ng/mL, which represents a decrease in neutralization activity by a factor of ∼7 (Table 1) compared to the Delta strain. For BQ.1.1 there is further decrease in Sotrovimab activity with an EC_50_ increasing to 1238 ng/ml, corresponding to a 12.6 fold decrease in neutralization activity when compared to Delta. For BQ.1.1 a greater loss of Sotrovimab activity has been recently reported using neutralization with pseudo-virus^10,24^, live virus^25^ and fusogenicity reporter assays^26^. The discrepancy between these different results may be explained by the fact that both studies using replicative virus used cell lines overexpressing ACE-2, which have been shown to artefactually underestimate the efficacy of S309 in *in vitro* tests^27^.

The neutralizing activity of Tixagevimab is very low against both BA.2 and BA.5 and is not restored in any other tested variants (Supp Fig.2, EC_50_ >5000 ng/L, see Table 1).

The other antibody of the Evusheld cocktail, Cilgavimab, which had regained neutralizing power against BA.2 and BA.5, completely lost its neutralizing activity against BQ.1 and BQ.1.1 (Supp Fig.2, EC_50_ >5000 ng/L, see Table 1). The same pattern is observed with both the XBB and BA.2.75.2 variants with no detectable neutralization. This loss of Cilgavimab activity directly affects the Evusheld cocktail with a loss of neutralizing activity against all new subvariants tested (Fig. 1, EC_50_ >10000 ng/L, see Table 1). These results are in line with recent studies aforementioned^10,24–26^.

Mechanistically, the loss of activity against the BA.5 subvariants BQ.1 and BQ.1.1 may be due to the to K444T mutation which is located in a region identified as critical for Cilgavimab neutralizing activity^2^. For BA.2.75.2 and XBB the loss of Cilgavimab activity could be due to the presence of two critical mutations: (i) G446S, responsible for the decrease of Cilgavimab activity against BA.1^22^, and (ii) R346T, which, in association with G446S, is responsible for a greater loss of Cilgavimab activity against BA1.1^28^ Finally, Bebtelovimab, the most recently produced antibody which has retained a strong activity against the Omicron BA.2 and BA.5 variants, also kept its activity against BA.2.75.2. However, it has lost all neutralizing activity against BQ.1, BQ.1.1 and XBB. This is likely due to the presence of K444T in BQ.1 and BQ.1.1 and V445P in XBB. These two residue positions have been determined by deep mutational scanning to be the major sites of mutational escape for Bebtelovimab^29^.

Altogether, we conclude that Sotrovimab retains some neutralizing activity against BA.2.75.2, XBB, BQ.1 and BQ.1.1 despite a further decrease in its activity compared to the BA.2 and BA.5 progenitors. This decrease remains limited but should be closely monitored as its clinical impact remains to be documented. Further investigation *in vivo* should be performed to ensure that the recommended dose of 500 mg is sufficient to provide the best expected therapeutic benefit. Regarding the Evusheld cocktail, there is a loss of neutralizing activity and it is likely that its *in vivo* activity could be jeopardized against these Omicron subvariants. These results illustrate the need for a strategy that offers a combination of antiviral molecules and therapeutic antibodies that offer a broader spectrum of activity or that effectively accompany the antigenic evolution of SARS-CoV-2.

## Acknowledgments

We thank Pr B.Lina for providing the SARS-CoV-2 BA.2, BA.5 and XBB strains. We thank Rayane Amaral and Camille Placidi-Italia for the technical help regarding the molecular biology experiments and the cell culture. We thank the Dr Cecile Baronti for her help and expertise regarding BSL3 experiments at UVE. We thank the staff of the National Reference Center for technical help, the team of the Mutualized Platform of Microbiology (P2M) at Institut Pasteur for help with sequencing and bioinformaticians of the Bioinformatics Hub at Institut Pasteur for help with sequence analysis. We are indebted to Dr. H. Mouquet (Institut Pasteur, Paris, France) for providing the Bebtelovimab/LY-Cov1404 antibody. We thank the public and private laboratories for providing specimens and aknowledge the EMERGEN genomic surveillance consortium. We acknowledge the authors, originating and submitting laboratories of the sequences from GISAID and GenBank. This work was performed in the framework of the Preclinical Study Group of the French agency for emerging infectious diseases (ANRS-MIE). It was supported by the ANRS-MIE (BIOVAR and PRI projects of the EMERGEN research program) and by the European Commission (European Virus Archive Global project (EVA GLOBAL, grant agreement No 871029) of the Horizon 2020 research and innovation program). The SVDW and ESL laboratories acknowledge funding from the European Commission (RECOVER project, grant agreement N° 101003589) of the Horizon 2020 research and innovation program and by the French Government’s Investissement d’Avenir program, Laboratoire d’Excellence “Integrative Biology of Emerging Infectious Diseases” (grant n°ANR-10-LABX-62-IBEID).The SVDW lab acknowledges funding from Santé publique France (the French national public health agency), the “Enhancing Whole Genome Sequencing (WGS) and/or Reverse Transcription Polymerase Chain Reaction (RT-PCR) national infrastructures and capacities to respond to the COVID-19 pandemic in the European Union and European Economic Area” Grant Agreement ECDC/HERA/2021/007 ECD. 12221, and the Caisse nationale d’assurance maladie (Cnam), the national health insurance funds. The ESL laboratory acknowledges funding from the INCEPTION program (Investissements d’Avenir grant ANR-16-CONV-0005), the NIH PICREID program (Award Number U01AI151758).

## Author contributions

FT, ESL and XDL proposed the study. FT, ESL, SVDW JC, FA and XDL designed and conceived the experiments. FT, EG, JTR and JC performed the experiments. FD produced and analyzed some of the viruses. JB, FL and ESL performed the phylogenetic analysis. FT, EG and JC analyzed the results. FT and XDL wrote the paper. FT, EG, ESL, SVDW, FA and XDL reviewed and edited the paper with input from all authors.

## Inclusion and diversity

We support inclusive, diverse and equitable conduct of research

## Star Protocol

### Key resources table

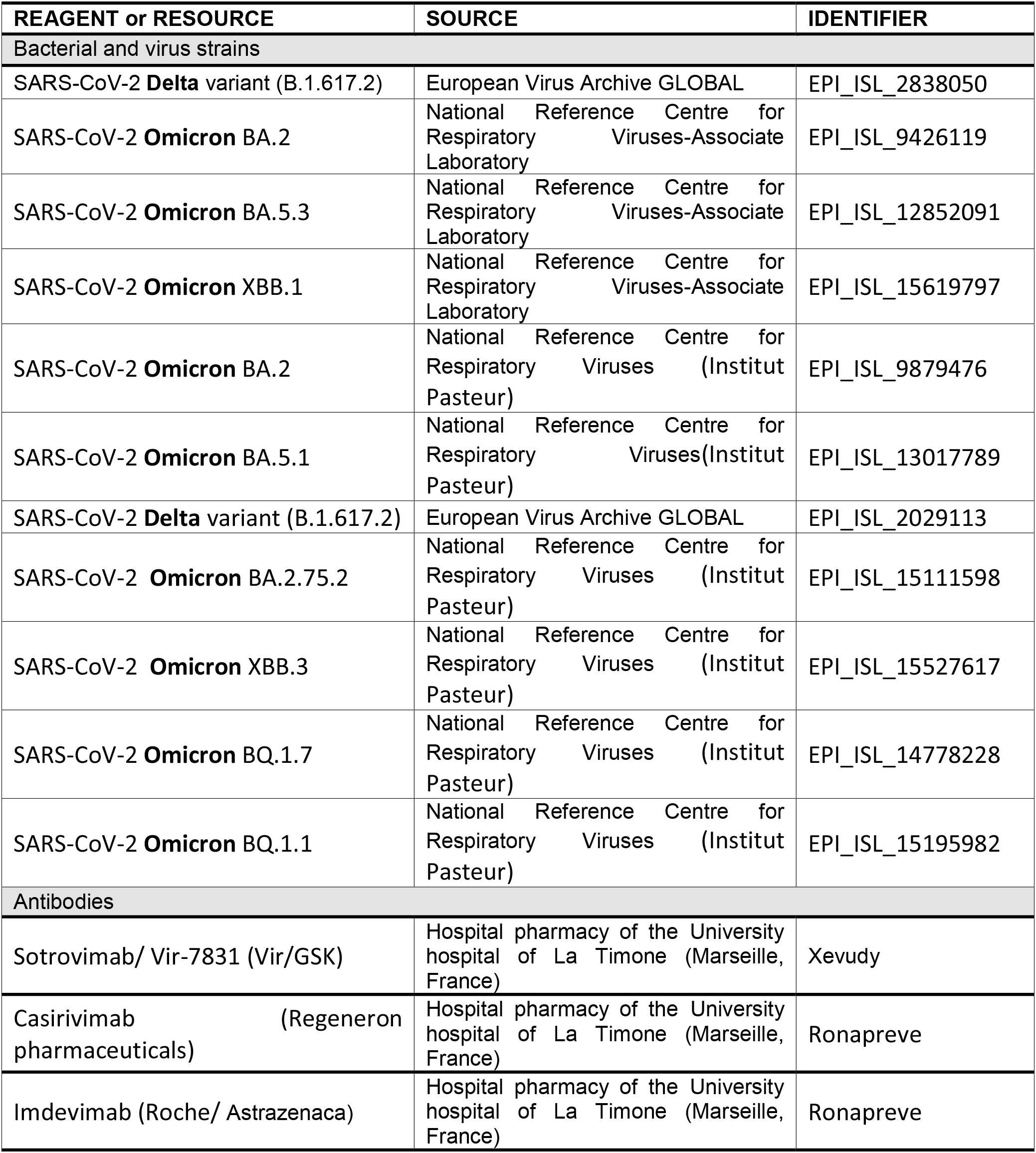
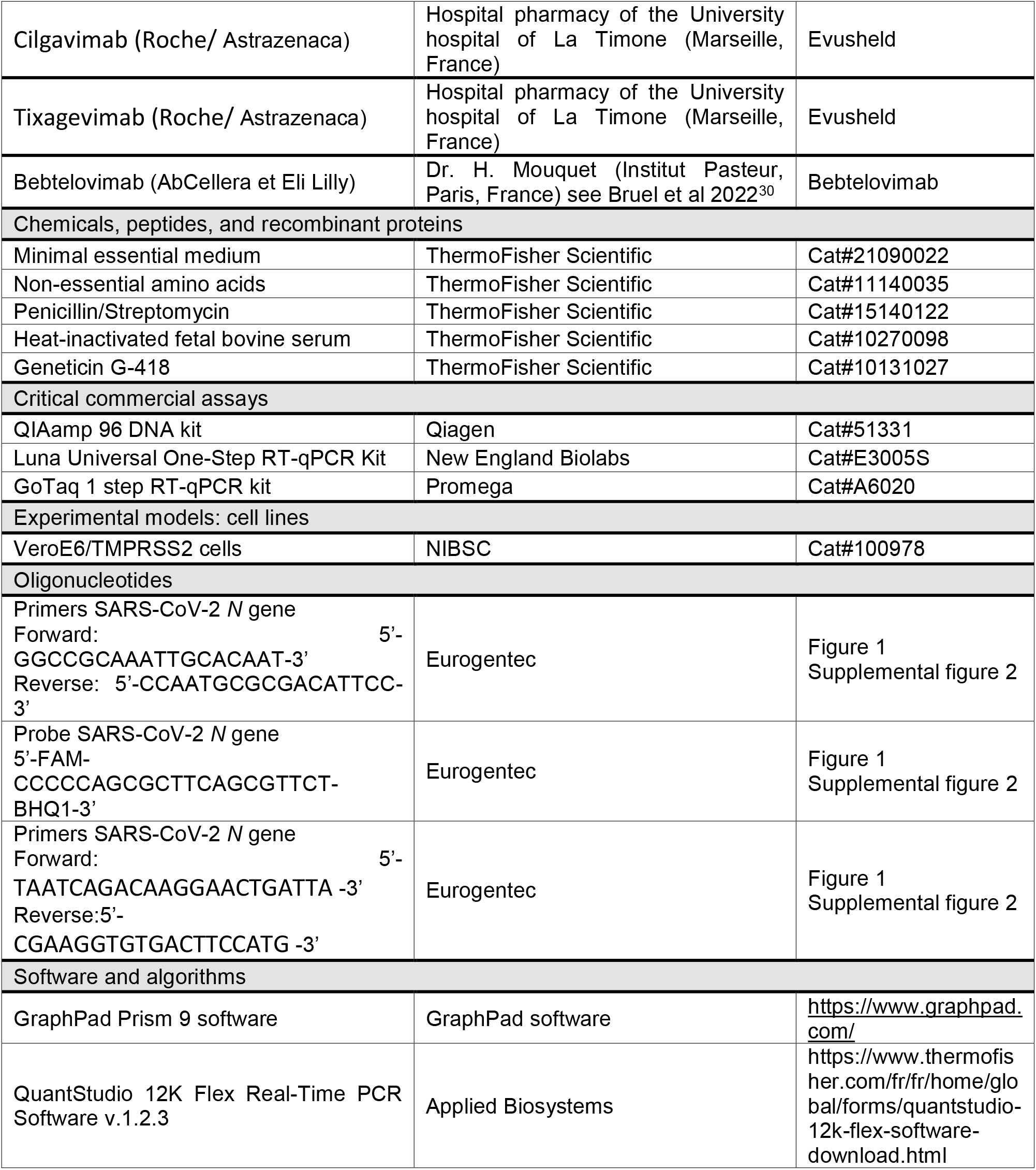

## Resource availability

### Lead contact

Further information and requests for resources and reagents should be directed to and will be fulfilled by the lead contact, Franck Touret.

### Materials availability

This study did not generate new unique reagents.

### Data and code availability

This study did not generate/analyze [datasets/code].

## Experimental model

### Cell line

VeroE6/TMPRSS2 cells (ID 100978) were obtained from CFAR and were grown in MEM (Minimal Essential Medium-Life Technologies) with 7 .5% heat-inactivated Fetal Calf Serum (FCS; Life Technologies with 1% penicillin/streptomycin PS, 5000U.mL^−1^ and 5000µg.mL^−1^ respectively (Life Technologies) and supplemented with 1 % non-essential amino acids (Life Technologies) and G-418 (Life Technologies), at 37°C with 5% CO_2_.

### Antibodies

Sotrovimab/ Vir-7831 was provided by GSK (GlaxoSmithKline). Casirivimab and Imdevimab (Regeneron pharmaceuticals), Cilgavimab and Tixagevimab (AstraZeneca) were obtained from the hospital pharmacy of the University hospital of La Timone (Marseille, France).

Bebtelovimab/LY-Cov1404 (Eli Lilly and Company) was kindly provided by Dr. H. Mouquet (Institut Pasteur, Paris) and its production and purification from Freestyle 293-F suspension cells was already described here^30^.

### Virus isolates

#### Isolates specific to UVE-Marseille

SARS-CoV-2 **Delta** variant (B.1.617.2) was isolated in May 2021 in Marseille, France. The full genome sequence has been deposited on GISAID: EPI_ISL_2838050. The strain, 2021/FR/0610, is available through EVA GLOBAL (www.european-virus-archive.com, ref: 001V-04282)

SARS-CoV-2 **Omicron** BA.2 strain hCoV-19/France/NAQ-HCL022005338701/2022 was obtained from Pr. B. Lina and the sequence is available on GISAID : EPI_ISL_9426119.

SARS-CoV-2 **Omicron** BA.5.3 strain hCoV-19/France/ARA-HCL022074071401/2022 was obtained from Pr. B. Lina and the sequence is available on GISAID : EPI_ISL_12852091.

SARS-CoV-2 **Omicron** XBB.1 strain hCoV-19/France/PAC-HCL022171892001/2022 was obtained from Pr. B. Lina and the sequence is available on GISAID : EPI_ISL_15619797.

#### Isolates specific to Institut Pasteur-Paris

SARS-Cov-2 B.1.617.2 (delta) strain hCoV-19/France/HDF-IPP11602i/2021 was supplied by the National Reference Centre for Respiratory Viruses hosted by Institut Pasteur (Paris, France) and headed by Pr. Sylvie van der Werf. The human sample from which strain hCoV-19/France/HDF-IPP11602i/2021 was isolated has been provided by Dr Guiheneuf Raphaël, CH Simone Veil, Beauvais France. Moreover, the strain hCoV-19/France/HDF-IPP11602i/2021 was supplied through the European Virus Archive goes Global (Evag) platform, a project that has received funding from the European Union’s Horizon 2020 research and innovation programme under grant agreement No 653316.

SARS-CoV-2 **Omicron** BA.2 (hCoV-19/France/PDL-IPP08031/2022) was isolated by the National Reference Center for Respiratory Viruses hosted by Institut Pasteur (Paris, France) and headed by Pr. Sylvie van der Werf, from a specimen provided by Dr. Bénédicte Lureau, CH Fontenay-le-Comte, 85201 Fontenay-le-Comte, France. The sequence is available on GISAID : EPI_ISL_9879476.

SARS-CoV-2 **Omicron** BA.5.1 (hCoV-19/France/BRE-IPP34319/2022) was isolated by the National Reference Center for Respiratory Viruses hosted by Institut Pasteur (Paris, France) and headed by Pr. Sylvie van der Werf, from a specimen provided by Dr Franck Kerdavid, Laboratoire d’Analyses Médicales, Alliance Anabio, 35520 Melesse, France. The sequence is available on GISAID : EPI_ISL_13017789.

SARS-CoV-2 **Omicron** XBB.3 (hCoV-19/France/HDF-IPP53307/2022) was isolated by the National Reference Center for Respiratory Viruses hosted by Institut Pasteur (Paris, France) and headed by Pr. Sylvie van der Werf, from a specimen provided by Dr Anne Vachée, CH de Roubaix, 59100 Roubaix, France. The sequence is available on GISAID : EPI_ISL_15527617.

#### Isolates common to UVE-Marseille and Institut Pasteur-Paris

SARS-CoV-2 **Omicron** BA.2.75.2 (hCoV19/France/IDF-IPP50347/2022) was isolated by the National Reference Center for Respiratory Viruses hosted by Institut Pasteur (Paris, France) and headed by Pr. Sylvie van der Werf, from a specimen provided by Dr Laura Djamdjian, CH de Gonesse, 95500 Gonesse, France. The sequence is available on GISAID : EPI_ISL_15111598

SARS-CoV-2 **Omicron** BQ.1.7 (hCoV-19/France/HDF-IPP49210/2022) was isolated by the National Reference Center for Respiratory Viruses hosted by Institut Pasteur (Paris, France) and headed by Pr. Sylvie van der Werf, from a specimen provided by Dr Arnaud Serpentini, Unilabs Henin Beaumont, 62110, Henin Beaumont, France. The sequence is available on GISAID : EPI_ISL_14778228

SARS-CoV-2 **Omicron** BQ.1.1 (hCoV-19/France/IDF-IPP50823/2022) was isolated by the National Reference Center for Respiratory Viruses hosted by Institut Pasteur (Paris, France) and headed by Pr. Sylvie van der Werf, from a specimen provided by Dr Beate Heym, Laboratoire des Centres de Santé et d’Hôpitaux d’IDF, 75020 Paris, France. The sequence is available on GISAID : EPI_ISL_15195982

All viral stocks were prepared by propagation in Vero E6 TMPRSS2 cells or in Vero E6 cells in the presence of TPCK trypsin for the BA.2, BA5.1 viruses used at Institut Pasteur.

All experiments involving live SARS-CoV-2 were performed in Biosafety Level 3 (BSL-3).

## Method details

### EC_50_ determination at UVE-Marseille

#### 1. Experiment conduction

a. One day prior to infection, 5×10^4^ VeroE6/TMPRSS2 cells per well were seeded in 100µL assay medium (containing 2.5% FBS) in 96 well culture plates.
b. The next day, antibodies were diluted in PBS with eleven ½ dilutions from 5000 to 4.8 ng/ml for Sotrovimab, Cilgavimab, Tixagevimab, Casirivimab and Imdevimab and its combination Evusheld. Then 25µL/well of the serial dilutions of antibodies were added to the cells in triplicate. Then, 25µL/well of a virus mix diluted in medium was added to the wells. Each well was inoculated with 100 TCID_50_ of virus which correspond here to a MOI at 0.002 as classically used for SARS-CoV-2^31^. Prior to the assay it was verified for each variant that with this MOI, viruses in the cell culture supernatants were harvested during the logarithmic growth phase of viral replication at 48 hours post infection ^21,22^. Four virus control wells were supplemented with 25µL of assay medium.

#### 2. Experimental analysis

Quantification of the viral genome by real-time RT-qPCR as previously described ^32^.

a. Nucleic acid from 100µL of cell supernatant were extracted using QIAamp 96 DNA kit and Qiacube HT robot (both from Qiagen).
b. Viral RNA was quantified by real-time RT-qPCR (GoTaq 1 step RT-qPCR kit, Promega). Quantification was provided by serial dilutions of an appropriate T7-generated synthetic RNA standard. RT-qPCR reactions were performed on QuantStudio 12K Flex Real-Time PCR System (Applied Biosystems) and analyzed using QuantStudio 12K Flex Applied Biosystems software v1.2.3. Primers and probe sequences, which target SARS-CoV-2 N gene, were: Fw: 5’-GGCCGCAAATTGCACAAT-3’ ; Rev : 5’-CCAATGCGCGACATTCC-3’; Probe: 5’-FAM-CCCCCAGCGCTTCAGCGTTCT-BHQ1-3’.

#### 3. Interpretation of the results

Viral inhibition was calculated as follow: 100* (quantity mean VC-sample quantity)/ quantity mean VC. The 50% effective concentrations (EC50 compound concentration required to inhibit viral RNA replication by 50%) were determined using logarithmic interpolation after performing a nonlinear regression (log(agonist) vs. response --Variable slope (four parameters)) as previously described ^21–23^. All data obtained were analyzed using GraphPad Prism 9 software (Graphpad software).

### EC_50_ determination at Institut Pasteur-Paris

#### 1. Experiment conduction

a. One day prior to infection, 3 × 10^3^ VeroE6/TMPRSS2 cells per well were seeded in 100µL assay medium (containing 2.5% FBS) in Black with clear bottom 384-well plates.
b. individual antibodies were added at indicated concentrations 2 h prior to infection. PBS (2μl) and remdesivir (25 µM; SelleckChem) controls were added in each plate. After the pre-incubation period, the virus inoculum (MOI 0.05 to 0.5 PFU/cell depending on the viral strains) was added to the cells. Following a one-hour adsorption at 37 °C, the supernatant was removed and replaced with 2% FBS/DMEM medium containing the individual antibodies at the indicated concentrations. Cells were incubated at 37 °C for 2 days.
c. Supernatants were harvested and heat-inactivated at 80 °C for 20 min.

#### 2. Experimental analysis

Detection of viral genomes from heat-inactivated samples was performed by RT-qPCR using the Luna Universal One-Step RT-qPCR Kit (New England Biolabs) with SARS-CoV-2 specific primers targeting the N gene region (5′-TAATCAGACAAGGAACTGATTA-3′ and 5′-CGAAGGTGTGACTTCCATG-3′) and with the following cycling conditions: 55 °C for 10 min, 95 °C for 1 min, for 1 cycle, followed by 95 °C for 10 s, 60 °C for 1 min, for 40 cycles in an Applied Biosystems QuantStudio 6 thermocycler.

#### 3. Interpretation of the results

Curve fits and IC50 values were obtained in Prism.

### Phylogenetic analysis

The full sequence dataset was downloaded from GISAID on Nov. 7^th^ 2022, and annotated using pangolin 4.1.3 and pangolin data v1.16^33^. The dataset was divided in 3 categories: 1) Sequences annotated as BQ.1.1 (7,810 sequences), BA.2.75 (5,491 sequences) and XBB (25 sequences); 2) sequences of interest (lineages close to BQ.1.1 or BA.2.75), annotated as B.1.1.529, BA.2, BE.1, BE.1.1, BE.1.1.1, and BQ.1 (1,047,744 sequences); and 3) all the other sequences (12,971,674 contextual sequences). We then used these categories to build a subsampled dataset using augur 5.0.1, with 25 XBB, 78 BQ.1.1 and 14 BA.2.75 (category 1), 21 sequences of interest (category 2), 250 contextual sequences (category 3), and 45 (resp. 9) additional sequences with genetic proximity to BQ.1.1 (resp BA.2.75). A total of 317 sequences (EPI_SET ID : EPI_SET_221207ws) were then analyzed using augur workflow 5.0.1^34^, and the annotated phylogenetic tree was converted to nexus using gotree v0.4.4a^35^.

