## Supplemental figures for "Enhanced neutralization escape to therapeutic monoclonal antibodies by SARS-CoV-2 Omicron sub-lineages"

[#] Co first authors
* Co-supervision

✉Correspondance : (FT), (XDL)

Keywords: SARS-CoV-2; Omicron; therapeutic monoclonal antibody; live virus

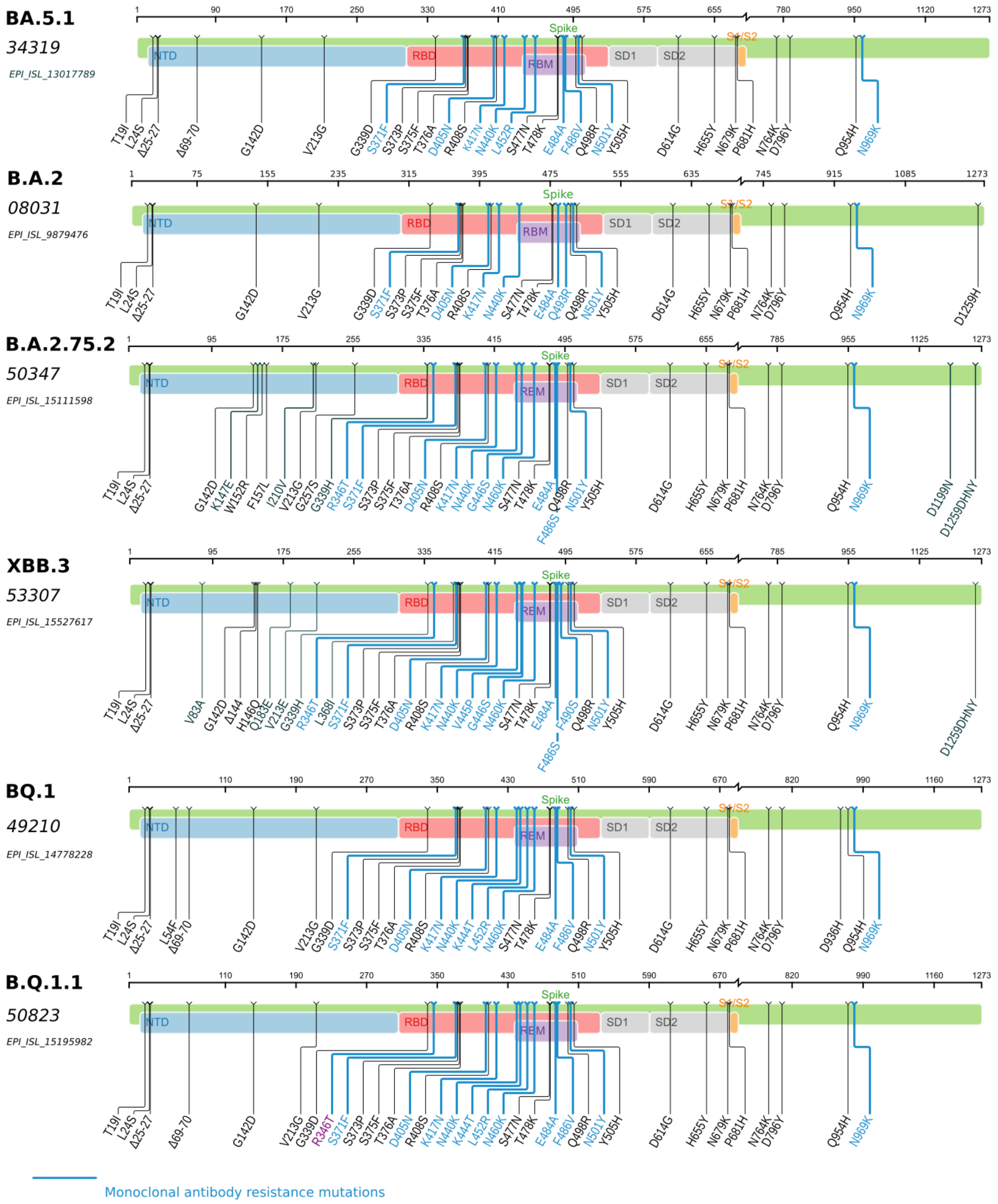


Supplemental figure1: Representation of mutations in Spike protein, for the main lineages studied (BA.5.1, BA.2, BA.2.75.2, XBB.3, BQ.1, and BQ.1.1), in comparison to Wuhan/hu-1/2019. Mutations conferring resistance to monoclonal antibodies are displayed in blue. These graphs have been created with the Sierra tool of the Stanford covdb database[1]

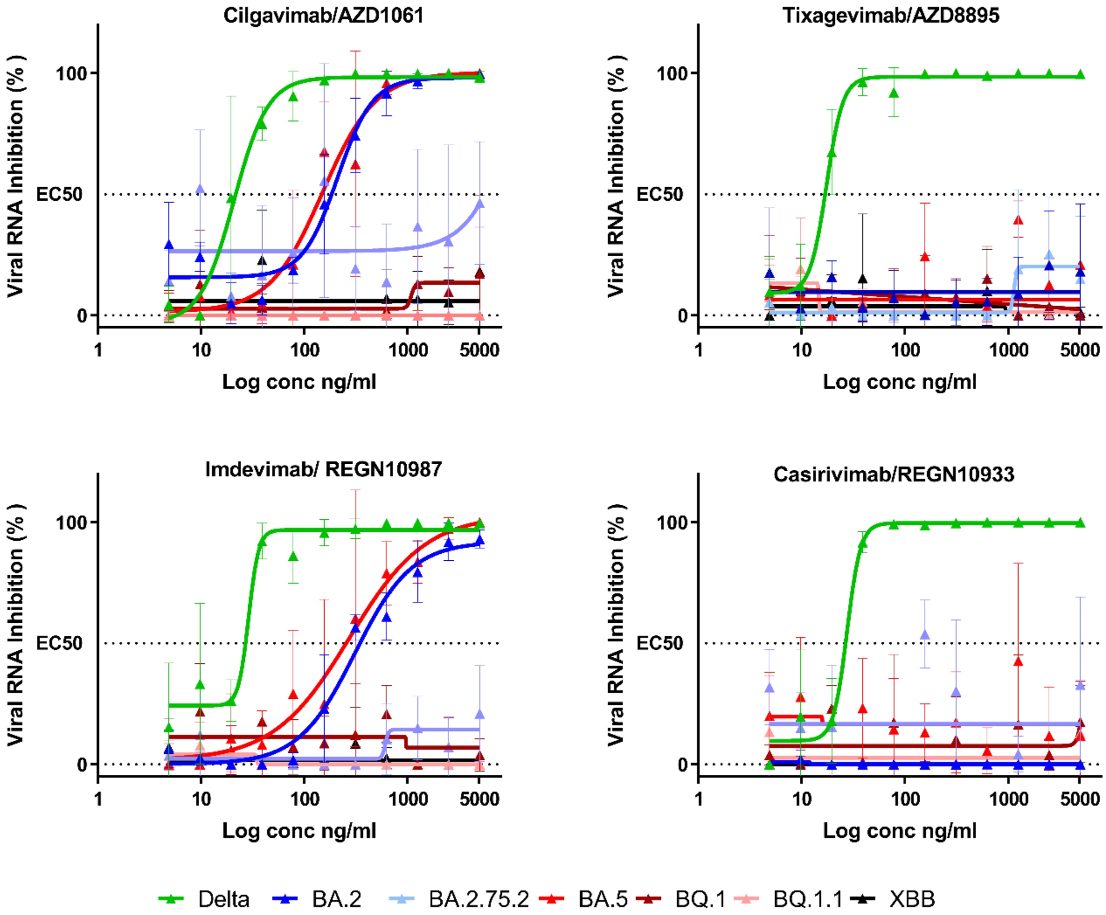


Supplemental figure 2: Dose response curves reporting the susceptibility of the SARS-CoV-2 Delta pre-omicron strain and Omicron subvariants to a panel of therapeutic monoclonal antibodies. Antibodies tested: Casirivimab/REGN10933, Imdevimab/REGN10987Tixagevimab/AZD8895, Cilgavimab/AZD1061. Data presented are from three technical replicates in VeroE6-TMPRSS2 cells, and error bars show mean±s.d.

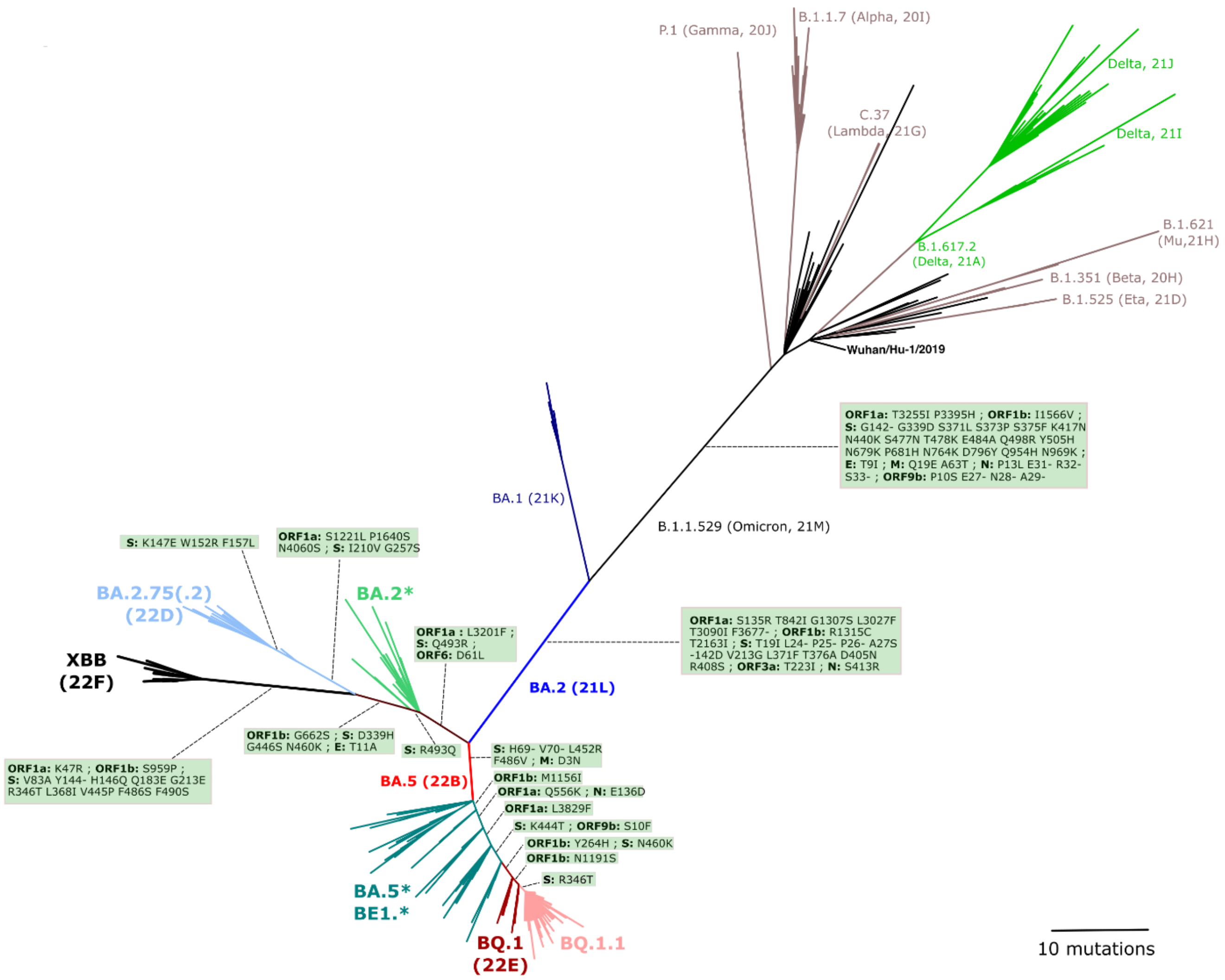


Supplemental figure 3: *Unrooted phylogenetic tree displaying BQ.1.1 and XBB lineages in the context of SARS-CoV-2 main lineages; amino-acid mutations are displayed on branches for the lineages of interest*.

1. Tzou, P.L., Tao, K., Pond, S.L.K., and Shafer, R.W. (2022). Coronavirus Resistance Database (CoV-RDB): SARS-CoV-2 susceptibility to monoclonal antibodies, convalescent plasma, and plasma from vaccinated persons. PLOS ONE *17*, e0261045. 10.1371/journal.pone.0261045.
